# SARS-CoV-2 infection in hiPSC-derived neurons is cathepsin-dependent and causes accumulation of HIF1ɑ and phosphorylated tau

**DOI:** 10.1101/2024.11.21.624622

**Authors:** Kettunen Pinja, Ruuska Janika, Quirin Tania, Ojha Ravi, Saber H Saber, Mohamed Shaker, Sean Morrison, Ernst Wolvetang, Merja Joensuu, Koistinaho Jari, Rolova Taisia, Balistreri Giuseppe

**Affiliations:** Helsinki Institute of Life Science, University of Helsinki, Helsinki, Finland; Department of Virology, Faculty of Medicine, University of Helsinki, Helsinki, Finland; Australian Institute for Bioengineering and Nanotechnology, The University of Queensland, Brisbane, Queensland 4072, Australia; Queensland Brain Institute, The University of Queensland, Brisbane, Australia; Zoology and Entomology Department, Faculty of Science, Assiut university, Assiut 71516, Egypt

**Keywords:** SARS-COV-2, entry mechanism, cathepsins, tau, neurodegeneration, HIF-1α

## Abstract

The severe acute respiratory syndrome coronavirus 2 (SARS-CoV-2) has been shown to infect the human brain and a subset of human neurons *in vitro*. We have previously demonstrated that the virus enters the human induced pluripotent stem cell (hiPSC)-derived neurons via an endosomal-lysosomal pathway, which is dependent on low levels of angiotensin-converting enzyme 2 (ACE2) and independent of transmembrane serine protease 2 (TMPRSS2). Here, we use hiPSC-derived neurons overexpressing ACE2 in co-culture with human astrocytes to show that the infection with both SARS-CoV-2 Wuhan and Omicron XBB.1.5 variants is dependent on cathepsins and can be efficiently blocked by an inhibitor of cathepsin B (CA-074-ME). The result was reproducible in non-transgenic hiPSC-derived cortical organoids. The cathepsin L inhibitor SB412515 was less effective against the Wuhan strain but equally effective against the Omicron variant. Using PCR and reinfection assays, we show that SARS-CoV-2 can replicate in neurons in 2D co-cultures. Interestingly, the infectivity of the newly produced virions declined at 24 hours post-infection despite a further increase in released viral RNA at later time points, suggesting the possible activation of an antiviral response in neurons and/or astrocytes, which is supported by a correspondent increase in the levels of secreted cytokines. Furthermore, the number of infected neurons decreased within five days, suggesting that SARS-CoV-2 infection eventually leads to the death of the target neuronal cell *in vitro*. The infection also caused the accumulation of the hypoxia-inducible stress factor HIF1-α in infected neurons under normoxia. Finally, we confirm and expand the previous finding that in SARS-CoV-2 infected neurons, the microtubule-associated protein tau is hyperphosphorylated at multiple loci, including S202/T205, and mislocalized to the soma of the infected neurons. Hyperphosphorylation and mislocalization of tau are hallmarks of Alzheimer’s disease (AD) and other ‘tauopathies’. Our data provides further evidence supporting the neurodegenerative potential of SARS-CoV-2 infection.

**Summary:** The recent COVID-19 pandemic has raised concerns about the potential for SARS-CoV-2 to infect the brain and worsen brain diseases like Alzheimer’s disease. Research has shown that SARS-CoV-2 can indeed infect the human brain, including a small number of neurons and other brain cells in laboratory settings.

In our previous studies, we identified the endosomal pathway as the route the virus uses to enter neuronal cells. In this study, we build on that work by demonstrating that inhibitors of endo-lysosomal cathepsin proteases can block this neuronal infection. We also found that infectious progeny virions are released from the infected neuronal cells.

Importantly, the infection proves harmful to the host cells, as evidenced by a decrease in the number of infected cells in experimental cultures over a five-day period. Additionally, we confirm and expand on earlier findings that SARS-CoV-2 infection leads to the phosphorylation and altered localization of the tau protein, a process associated with brain diseases like Alzheimer’s.

Finally, we observed an increase in the production of inflammatory cytokines following neuronal infection with SARS-CoV-2, along with an accumulation of the stress marker protein HIF-1α in neurons. This protein has been linked to other viral infections and Alzheimer’s disease. Overall, our data suggest that SARS-CoV-2 exhibits neurodegenerative characteristics.

## Introduction

SARS-CoV-2 is a respiratory coronavirus responsible for the global COVID-19 pandemic. It mainly targets epithelial cells of the respiratory tract, but it can also infect other organs, including the brain. Due to the high prevalence of neurological symptoms related to SARS-CoV-2 infection (1,2), interest in the possible neuronal tropism of SARS-CoV-2 has increased.

Since the beginning of the pandemic, post-mortem studies have demonstrated the presence of viral components, such as proteins and RNA, in the brain tissues of individuals who have succumbed to SARS-CoV-2 infection (3–6). Anatomical changes, such as a reduction in grey matter thickness and global brain volume, are also detectable in the brains of COVID-19 patients (7). Stem cell-based *in vitro* models have shown that the virus is capable of infecting brain tissues, including the epithelial cells of the choroid plexus (8,9), as well as a small subset of neurons and astrocytes (6,10–13), and possibly even microglia (14). Moreover, post-mortem and cell culture studies have shown that SARS-CoV-2 infection can induce changes at the cellular level associated with Alzheimer’s disease (AD). These changes include altered neuronal morphology (12,15), and survival (6,10,16), microglial activation (17,18), NLRP3 inflammasome activation (17–19), cytokine production (10,17,20,21), astrogliosis (6,13,20), and microglia-mediated synapse elimination (16). Thus, investigating the mechanisms of SARS-CoV-2 entry into brain cells and identifying the key host factors involved in this process is crucial for developing strategies to prevent or reduce the potential spread of the virus in brain tissue.

There are two main routes for SARS-CoV-2 to productively infect cells: A) through direct fusion of the viral membrane or early endosomes where the viral spike protein is cleaved by surface proteases such as TMPRSS2 or membrane-type matrix metalloproteinases (22), leading to the release of the viral genome into the cytoplasm, or B) following receptor-mediated endocytosis and cleavage of the viral spikes in endo-lysosomes by proteases such as cathepsins (23) Cathepsins are pH-dependent endo-lysosomal proteases involved in protein degradation, metabolism, and host immune function. They are divided into three families: serine proteases (cathepsin A and G), aspartic proteases (D and E), and cysteine proteases (B, C, F, H, K, L, O, S, V, X, and W) (24). Based on computational models, multiple cathepsins can cleave the SARS-CoV-2 spike protein (25). Among these, cathepsin L has received the most attention as the mediator of SARS-COV-2 host cell entry in cells devoid of cell surface serine proteases (26,27).

We have recently demonstrated that SARS-CoV-2 enters human induced pluripotent stem cell (hiPSC)-derived neurons through the endo-lysosomal compartment in an ACE2-dependent and TMPRSS2-independent manner (12). However, we did not identify the endo-lysosomal proteases needed for neuronal SARS-CoV-2 infection.

Additionally, the low infection rate made it difficult to confirm whether SARS-CoV-2-infected neurons produced infectious virions. This was likely due to the low expression of surface proteins such as ACE2 (12,28,29).

To investigate the early phases of the virus life cycle, we used ACE2-overexpressing hiPSC neurons to demonstrate that SARS-CoV-2 infection is productive and can be blocked by the cathepsin B inhibitor CA074-ME. The results were confirmed using non-transgenic hiPSC-derived cerebral organoids containing choroid plexus-like cells (30,31). We also show that the infection causes pathological changes in the infected neurons, such as increased phosphorylation of the microtubule-associated protein tau at multiple sites as well as its mislocalization – a hallmark of the late-onset AD and other neurodegenerative diseases commonly known as ‘tauopathies’ (32).

## Results

### Characterization of ACE2-overexpressing hiPSC-derived neurons

Using a 2D neuron-astrocyte co-culture model, we have previously shown that only a small subset of hiPSC-derived mature neurons is susceptible to SARS-CoV-2 infection (12). The infection was inhibited by antibodies against ACE2 and inhibitors of early-to-late endosome transport (i.e., the PIKfive inhibitor apilimod dimesylate), but not by inhibitors of serine proteases such as TMPRSS2 (i.e. nafamostat mesylate)(12), consistent with undetectable mRNA levels of this protease as measured by qRT-PCR (12). Although this model system confirmed SARS-CoV-2’s ability to infect and replicate in a subset of human neurons, the limited infectivity made further investigations challenging.

Having established that SARS-CoV-2 infection of hiPSC-derived neurons requires ACE2 and endosomal transport, we set out to identify the endo-lysosomal proteases responsible for the cleavage of the viral spike protein, which allows membrane fusion and release of the viral genome into the host cytosol.

To this end, we created a hiPSC-derived neuron-astrocyte co-culture model where the neurons stably overexpress human ACE2 (Fig. 1A; All the original data shown in the graphs presented in this work are available in supplementary data file “S1 Data”). Although transduction with an ACE2-expressing lentivirus induced robust expression of ACE2 at mRNA level in lung carcinoma cell line A549, it induced only a modest increase in ACE2 in hiPSCs and 35-day-old neuron-astrocyte co-cultures (Fig. 1B).

**Figure 1:**
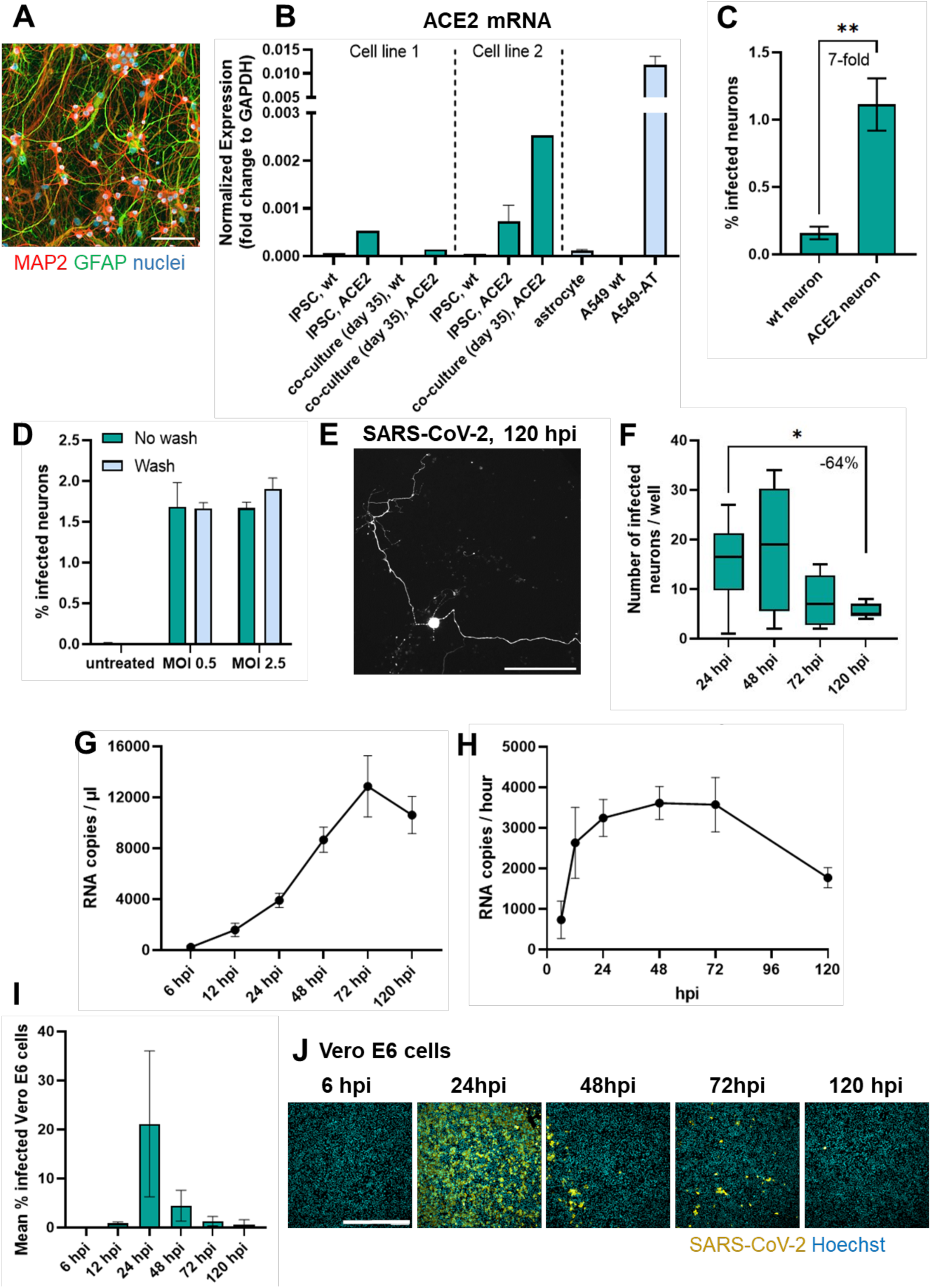
A. Representative image of a neuron-astrocyte co-culture. Scale bar 100 µm. MAP2 (microtubule-associated protein 2). GFAP (Glial fibrillary acidic protein). ACE2 mRNA expression in wildtype and ACE2-transduced hiPSCs, 35-day-old neuron-astrocyte co-cultures, astrocytes and negative and positive control cell lines (A549 wildtype and A549-ACE2-TMPRSS2 (AT)), respectively. Two hiPSC-lines derived from two individuals were used for brain cell differentiation. C. Comparison of the infection rate between co-cultures that contain control (wt neuron) or ACE2 overexpressing neurons (ACE2 neuron). Statistical significance was tested using Mann-Whitney U test. **, P ≤ 0.01. N = 6 replicate wells. D. The effect of viral MOI and washing on infection rate. E. A representative image of an infected neuron with intact neurites at 120 hpi. Scale bar 300 µm. F. Comparison of the number of infected neurons per 96-well at 24, 48, 72 and 120 hpi. Statistical significance tested using Kruskal-Wallis test with Dunn’s correction for multiple comparison. *, P ≤ 0.05. N = 3-14 replicate wells. G. Viral RNA copy number per one µL of conditioned medium at 6, 12, 24, 48, 72 and 120 hpi. N = 4 replicate wells. H. Production of viral RNA copies per hour at 6, 12, 24, 48, 72 and 120 hpi. I. Infectivity of susceptible Vero E6 cells by conditioned medium collected from infected neuron-astrocyte co-cultures at 6, 12, 24, 48, 72 and 120 hpi. N = 4 replicate wells. J. Representative images of Vero E6 cultures treated with conditioned medium from infected neuron- astrocyte co-cultures are 6, 24, 48, 72, and 120 hpi. Scale bar 800 µm. Only significant results are denoted.

Furthermore, the ACE2 expression varied between hiPSC lines from two different donors (Fig. 1B). Despite the modest increase in ACE2 mRNA levels, the infection rate of ACE2-transduced neurons increased 2-10 times compared to non-transduced controls (Fig. 1C). Increasing virus multiplicity of infection (MOI) or the washing of the cultures with fresh medium two hours after infection did not affect infectivity indicating that susceptible neurons were infected already at low MOI (Fig. 1D).

Immunostaining with cell type-specific antibodies indicated that the infected cells were almost exclusively neurons. Astrocyte infection was rarely observed (one infected cell per tens of thousands of cells). However, astrocyte infection was increased by ACE2 overexpression (Supplementary figure 1), suggesting that astrocytes can support SARS-CoV-2 infection and that the level of ACE2 expression is the limiting factor at least *in vitr*o.

Occasionally, we detected the cell-to-cell fusion of infected neurons (detected as multinucleated neurons stained with antibodies against the viral N protein) (Supplementary figure 2), which is consistent with earlier reports (33,34), as well as morphological changes (Supplementary figure 2) reminiscent of the SARS-CoV-2- induced cell membrane blebbing previously reported by Mignolet et. al. (35).

### Neuronal survival and morphology

To follow the impact of viral infection on neuronal survival and morphology, infected neurons were imaged after immunostaining at 24, 48, 72, and 120 hours post-infection (hpi) in neuron-astrocyte co-cultures. Roughly 90% of infected neurons had intact neurites at 24 hpi (89.5% ± 13.8%). The remaining infected neurons had a globular morphology, suggesting that they have retracted neurites. Even at 120 hpi, around 80% of the remaining infected neurons had their neurites intact (Fig. 1E), which indicates that infected neurons can persist in the culture for multiple days.

However, we detected a 64% reduction in the number of infected neurons between 24 and 120 hpi (Fig. 1F). Surprisingly, we were unable to detect signs of apoptosis in infected cells by TUNEL, propidium iodide, or anti-cleaved caspase-3 staining at any time point (Supplementary figure 2, D).

### Neuronal infection by SARS-CoV-2 is productive

To determine whether the neuronal infection was productive, we measured the release of viral RNA copies from SARS-CoV-2-treated neuron-astrocyte co-cultures using qRT-PCR. The results, shown as released viral RNA copies per µL (Fig. 1G) and viral RNA copies per hour (Fig. 1H) at 6-120 hpi, indicate that the release of viral RNA in the extracellular space increased until 72 hpi after which the kinetic of release slowed down (Fig. 1G, H).

To investigate whether the produced virions were capable of infecting new cells, we treated a susceptible cell line Vero E6 with conditioned medium collected from the SARS-CoV-2 infected neuron-astrocyte co-cultures at different time points. Our quantification indicated that the progeny virions released from infected neurons within 24 hpi were highly infectious (Fig. 1I, J). Interestingly, while the viral RNA copy number increased in neuronal media until 72 hpi (Fig. 1G, H), the infectivity of released virions peaked at 24 hpi and rapidly declined at later time points (Fig. 1I, J). Taken together, this data suggests that after 24 hpi the infected cells may mount an antiviral response, which does not block release but reduces the infectivity of the newly produced virions.

### Cathepsin inhibitors block SARS-CoV-2 infection in neurons

To investigate the involvement of different cathepsins in SARS-CoV-2 entry into neurons, we tested the efficacy of two well characterized cathepsin inhibitors, CA- 074-ME and SB412515, in inhibiting SARS-CoV-2 neuronal infection. Cathepsin inhibitor CA-074-ME has the highest affinity to cathepsin B (Ki = 2-5 nM compared to cathepsin H and L: Ki = 40 – 200 µM, MedChemExpress, New Jersey, USA).

SB412515 has the highest affinity to cathepsin L (IC50 = 0.85 nM compared to cathepsin B IC50 = 85.1 nM, Cayman Chemical, Michigan, USA).

Confirming our previous results obtained in cultures endogenously expressing ACE2 (36), in the ACE2-overexpressing model used here the PIKfyve inhibitor apilimod, which inhibits endosomal maturation and acidification, blocks 70% of the infection at 0.2 µM compared to DMSO. In contrast, and consistent with our previous results (36), 25 µM nafamostat, a broad-spectrum serine protease inhibitor, which blocks the TMPRSS2 cleavage of the SARS-CoV-2 spike protein at or near the cell surface, was not as effective (36% reduction compared to DMSO) (Fig 2A). Thus, as with our previously used non-transgenic co-culture models, SARS-CoV-2 infects ACE2- overexpressing neurons via the endo/lysosomal pathway.

**Figure 2:**
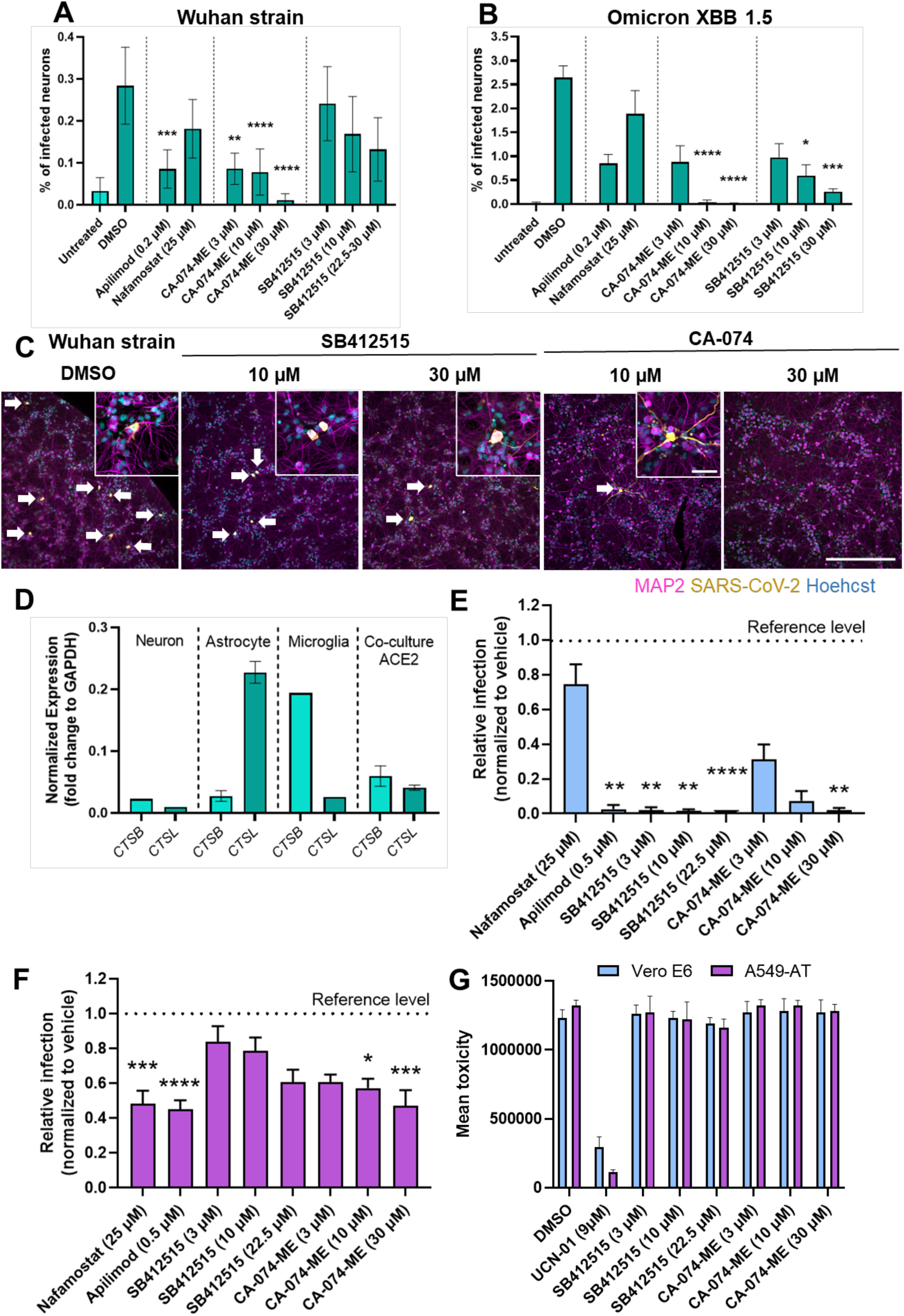
A. Effectiveness of 0.2 µM apilimod, 25 µM nafamostat, 3-30 µM CA-074- ME and 3-30 µM SB412515 in blocking neuronal infection by the Wuhan strain of SARS-CoV-2. For each drug treatment, values were normalized to the corresponding concentrations of vehicle (DMSO) controls. Analysis included 6 replicate wells in two independent batches. B. Effectiveness of 0.2 µM apilimod, 25 µM nafamostat, 3-30 µM CA-074-ME and 3-30 µM SB412515 in blocking neuronal infection by the Omicron XBB.1.5 strain of SARS-CoV-2. N = 6 replicate wells. C. Representative images of data shown at A. Scale bars 600 µm (main) and 100 µm (zoom-in). D. mRNA expression of cathepsin B (CTSB) and L (CTSL) in hiPSC-derived neurons, astrocytes, microglia, and neuron-astrocyte co-cultures. Data shown as fold change to the housekeeping gene *GAPDH*. E. Effectivity of 25 µM nafamostat, 0.5 µM apilimod, 3-22.5 µM SB41251 and 3-30 µM CA-074-ME in blocking SARS-CoV-2 infection in Vero E6 cells, F. and A549-AT cells. The data is displayed as fold-change to DMSO (reference level). N = 6 replicate wells. G. Cytotoxicity assay of 3-22.5 µM SB41251 and 3-30 µM CA-074-ME in Vero E6 and A549-AT cell lines. DMSO was used as negative and 9 µM UCN-01 as a positive control. N = 8 replicate wells. All statistical significances were assessed using Kruskal-Wallis test with Dunn’s correction for multiple comparison. All comparisons were made to the SARS-CoV-2 infected DMSO control. Only significant results are denoted. *, P ≤ 0.05; **, P ≤ 0.01; ***, P ≤ 0.001; ****, P ≤ 0.0001.

Results from five independent experiments showed that the cathepsin B inhibitor CA- 074-ME reduced infection by the Wuhan strain on average by 90% at 10µM (mean fold-change to DMSO: 0.10, 95% CI: 0.03 – 0.17, P = 0.01) and 88% at 30µM (mean fold-change to DMSO: 0.12, 95% CI: 0.00 – 0.38, P < 0.01). The cathepsin L inhibitor SB412515 was less effective against the Wuhan strain (31%, 54%, and 50% reductions at 3, 10 and 22.5 µM concentrations). Similar results were obtained when the data was pooled from two experiments using two different hiPSC lines as shown in Fig. 2A. Representative images of the cultures are shown in Figure 2C.

CA-074-ME also strongly reduced the infection by the Omicron XBB.1.5 strain (99% reduction at 10µM and 100% at 30µM as compared to DMSO; Fig 2B). SB412515 was more effective against the Omicron as compared to the Wuhan strain (78% reduction at 10µM and 90% at 30 µM as compared to DMSO; Fig. 2B).

The stronger effect of the cathepsin B inhibitor in neuronal cultures compared to the cathepsin L drug prompted us to measure the relative mRNA expression levels of the respective proteases in neurons and astrocytes. The two-fold higher neuronal expression levels of cathepsin B mRNA compared to cathepsin L (Fig. 2D) are consistent with the result that CA-074-ME was more effective against the Wuhan strain as compared to SB412515. In comparison, astrocytes expressed 23-fold higher levels of cathepsin L mRNA and microglia 8-fold higher levels of cathepsin B mRNA than neurons (Fig. 2D).

To evaluate the selectivity of the two drugs towards cathepsins in cells where the infection is known to require either cathepsin L or TMPRSS2 as controls, we tested the efficacy of CA-074-ME and SB412515 on two highly susceptible cell lines, Vero- E6 and A594-AT. Vero E6 cell line expresses ACE2 and cathepsin L but not TMPRSS2 (37). The A549-AT line, on the other hand, has been genetically engineered to overexpress both ACE2 and TMPRSS2 (38). The infection pathways for these cell lines are known. SARS-CoV-2 infection of Vero-E6 occurs in the endo- lysosomal compartment, which is confirmed here by the ability of apilimod and inability of nafamostat to block the infection (Fig. 2E, Vero E6, Sup.3). Both SB412515 and 30 μM CA-074-ME blocked infection in Vero-E6 cells (Fig. 2E, Sup.3). However, unlike the results obtained in neurons, SB412515 was effective even at the lowest concentration, which is consistent with an earlier report (39). In comparison, SARS-CoV-2 can infect the A594-AT line via both cathepsin and TMPRSS2 pathways, which is reflected here in the smaller effect of individual cathepsin inhibitor drugs (Fig. 2F, Sup.3). None of these drugs were cytotoxic at the concentrations and incubation times used in the study (Fig. 2G). Taken together, these data suggest that the used drugs displayed a sufficient specificity within the range of concentrations tested.

To validate our results in a different non-transgenic hiPSC-based model, we tested the cathepsin B inhibitor CA-074-ME and the two control drugs, apilimod and nafamostat, in choroid-plexus-containing cortical organoids (Fig. 3A). Neuronal infectivity by SARS-CoV-2 has been modest in many 3D organoid models (8,9,40). A recently described organoid model, which contains choroid plexus-like structures, supports a relatively high neuronal infection rate without the need for overexpression of ACE2 or other entry factors (31), and infection in neurons requires endosome transport (doi.org/10.1101/2023.03.03.530798). Thus, SARS-CoV-2 infection was more robust in these organoids compared to the 2D co-cultures (Fig. 3A).

**Figure 3.**
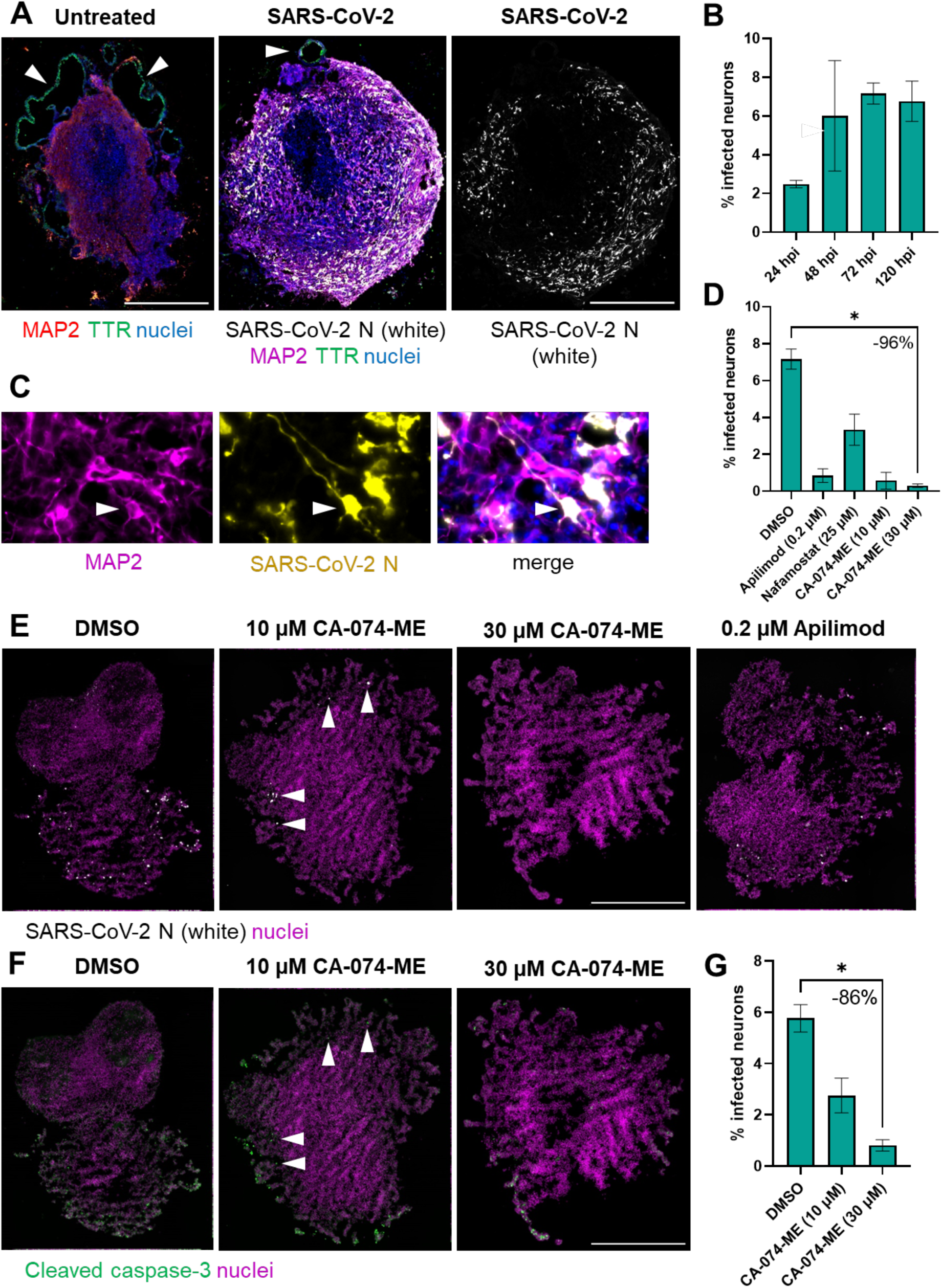
A. Representative images of untreated and SARS-CoV-2-infected cortical organoids with choroid plexus (arrowheads). Scale bars 300 µm. B. Quantification of the percentage of infected neurons at 24, 48, 72 and 120 hpi. C. Close-up of SARS- CoV-2 infected neurons (MAP2+). An infected cell marked with arrowheads. D. Quantification of the percentage of SARS-CoV-2 infected neurons in infected organoids with (10 and 30 µM CA-074-ME) and without (DMSO) drug-treatment at 72 hpi. E. Representative images of SARS-CoV-2 positive cells in infected organoids with (10 and 30 µM CA-074-ME) and without (DMSO) drug-treatment at 72 hpi. Arrowheads mark SARS-CoV-2 infected cells. Scale bar 500 µm. F. Representative images of cleaved caspase-3 positive neurons with and without drug treatment. Arrowheads correspond to figure. Scale bar 500 µm. D. G. Quantification of the % of cleaved caspase-3 positive neurons with and without drug treatment. All statistical significances were assessed using Kruskal-Wallis test with Dunn’s correction for multiple comparison. N = 3 organoids. Only significant results are denoted. *, P ≤ 0.05; **, P ≤ 0.01

Interestingly, the kinetics of virus spread in the 3D organoids were slower compared to 2D cultures with infection peaking at 48 hpi in 2D cultures (Fig. 1F) versus at 72 hpi in organoids (Fig. 3B). This could be due to a difference in the diffusion rate of viral particles in Matrigel-embedded organoids compared to 2D culture monolayers, and/or due to a difference in the kinetics of the virus reproduction and reinfection.

Consistent with the results presented by Shaker et. al. (31), the infected cells co- localized with MAP2-reactive fluorescent signal suggesting that the infected cells are mainly neurons (Fig. 3C). Treatment of the infected cultures with CA-074-ME decreased the percentage of infected MAP2-positive (MAP2+) cells in a dose-dependent manner, on average by 96% at 30 μM concentration compared to DMSO (Fig. 3E), indicating that the infection occurred mainly via endosomes and required cathepsins.

Consistently with our earlier report (30), apilimod reduced the percentage of infected MAP2+ cells on average by 88% at 0.2 μM concentration (Fig 3D, E), while high concentration of the TMPRSS2 inhibitor nafamostat was less effective (Fig. 3E).

A dose-dependent reduction in the percentage of the cleaved caspase-3 positive MAP2+ area in CA-074-ME-treated organoids shows that the drug did not induce apoptosis but rather decreased it (86% reduction at 30 µM concentration) compared to DMSO (Fig 3F, G), suggesting a neuroprotective rather than toxic effect.

Neuroprotective and antiapoptotic effects of cathepsin B inhibitors such as CA-074-ME have been previously described (41–43) independently of the SARS-CoV-2 infection.

### Tau phosphorylation increases in infected neurons

The prevalence of COVID-19-associated neurocognitive symptoms(1,2) has raised concern over the putative neurodegenerative potential of SARS-CoV-2 (44). To check for the signs of neurodegeneration in infected neurons, we immunostained 2D neuron-astrocyte co-cultures with an antibody specific for tau protein phosphorylated at serine 202 and threonine 205 (AT8) at different times during infection. We detected on average 1.5-fold increase in phosphorylated tau in the soma of infected neurons at 24, 48 and 72 hpi (Fig. 4A, B, C, D). Tau phosphorylation at serine 396, on the other hand, did not change significantly at 24 hpi (Fig. 4E, F). Hence, we focused on tau phosphorylation at Ser202/Thr205.

**Figure 4:**
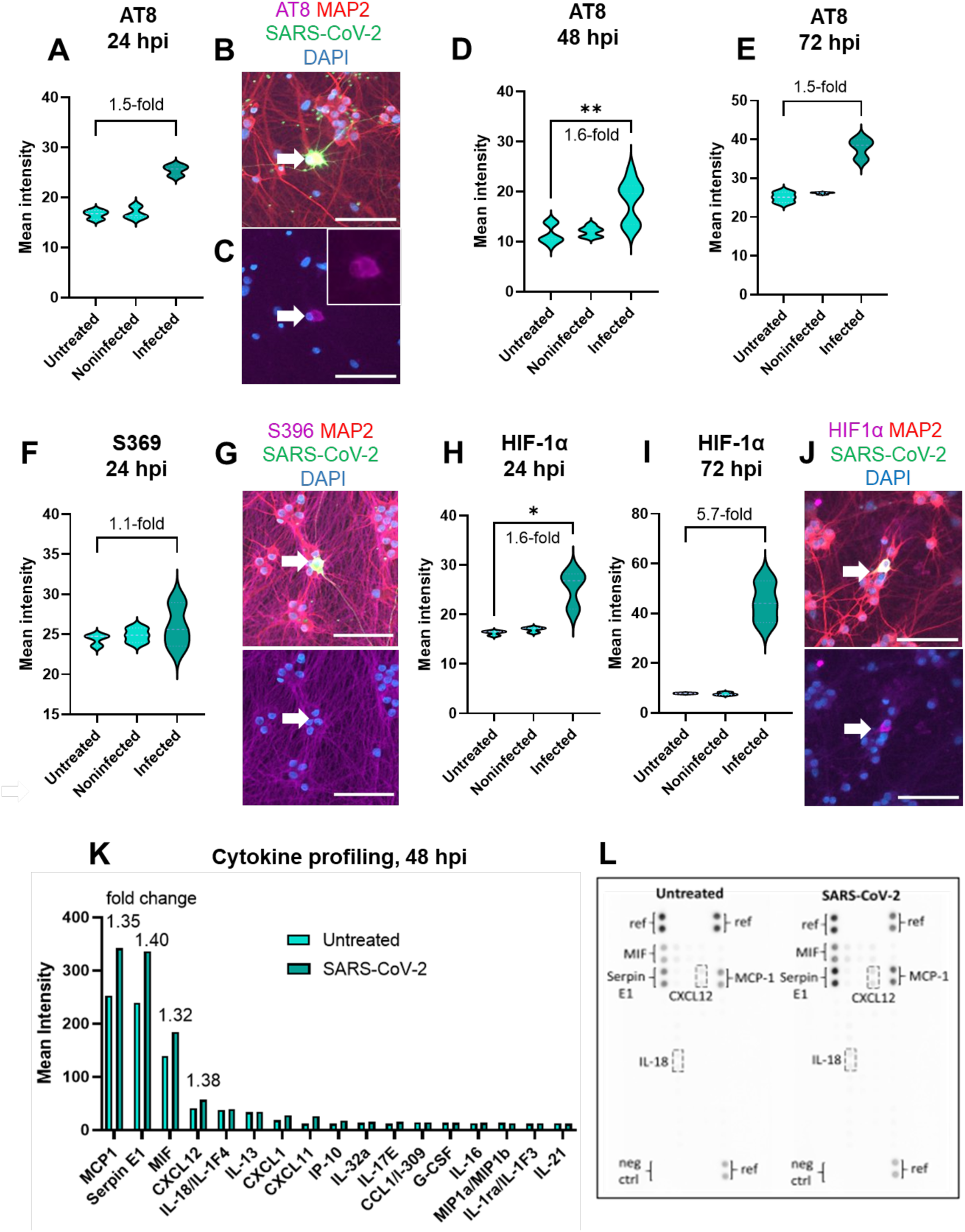
A, D, E. Mean intensity of AT8 antibody signal in untreated, treated but noninfected, and infected neurons at 24, 48 and 72 hpi. N = 2-6 replicate wells. B & Representative images of AT8 staining at 24 hpi. F. Mean intensity of anti-S396 signal in untreated, treated but noninfected, and infected neurons at 24 hpi. G. Representative images of anti-S396 staining at 24 hpi. N = 3 replicate wells. H & I. Mean intensity of anti-HIF-1α signal in untreated, treated but noninfected, and infected neurons at 24 and 72 hpi. N = 3 replicate wells in one batch. J. Representative images of anti-HIF-1α staining at 24 hpi. K. Secreted cytokine profile of SARS-CoV-2 infected thick neuron-astrocyte co-cultures and untreated controls at 48 hpi. Fold-changes to control are shown next to the four main hits. N = one sample pooled from two replicate wells. L. Annotated image of the membrane used to detect secreted cytokine profile of a SARS-CoV-2 infected neuron-astrocyte co-culture and positive control. All statistical significances were tested using Kruskal-Wallis test with Dunn’s correction for multiple testing. Only significant results are denoted. * ≤ 0.05, ** P ≤ 0.01. Scale bars: 100µm.

### Exposure to SARS-CoV-2 induces the stabilization of the stress factor HIF-1ɑ in neurons and increased production of cytokines

An upregulation of hypoxia-inducible factor 1-ɑ (HIF-1ɑ) has been reported in AD (45). Under physiological conditions (i.e., normoxia and absence of stressors), HIF- 1ɑ undergoes rapid turnover. Hypoxia and a range of cellular stressors stabilize the transcription factor, activating stress-sensitive transcriptional responses, including antiviral responses, that can lead to cell death or survival, depending on the intensity and duration of the stress. We detected an increase in HIF-1ɑ immunofluorescence in the soma of infected neurons. The level of HIF-1ɑ increased by 1.6-fold at 24 hpi (Fig. 4G, H) and by 5.7-fold at 72 hpi (Fig. 4I, J). This result suggests that HIF-1ɑ is stabilized by the presence of SARS-CoV-2 in hiPSC neurons.

The production of cytokines and chemokines promotes an anti-viral innate immune response (46,47). Using a cytokine profiler array comprising of 36 human cytokines and chemokines (Sup. 4A), we screened for cytokines and chemokines secreted by neuron-astrocyte co-cultures and detected increased levels of macrophage migration inhibitory factor (MIF), monocyte chemoattractant protein-1 (MCP-1/CCL2), Serpin E1 and CXC motif chemokine ligand 12 (CXCL12) in response to SARS-CoV-2 (Fig. 4K,L). We saw no marked difference in interferon gamma-induced protein 10 (IP- 10/CXCL10) levels between infected cultures and controls using the profiler or a cytokine bead array (Supplementary figure 4). We also tested the secretion of interleukin 6 and 8 (IL-6 and IL-8) from neuron-astrocyte co-cultures, but they were not detected in the cell culture medium.

### The effect of the presence of microglia on neuronal infection rate

Next, we addressed the effect of microglia on neuronal SARS-CoV-2 infection by establishing triplecultures containing neurons, astrocytes, and microglia. We had to modify the composition of cell culture medium by adding microglial growth and survival promoting factor IL-34 and by removing growth factor CNTF, the presence of which induced a hyperactivation of microglia and the loss of the co-culture. The obtained co-cultures exhibited around 28% ± 4% microglia (an average of three untreated and three infected wells) with elongated morphology in close contact with neural projections (Fig. 5A, B, C). A representative image of a SARS-CoV-2-infected neuron in microglia-containing triplecultures is shown in Fig. 5D. We did not detect microglial infection by SARS-CoV-2 Wuhan or Omicron XBB.1.5 strains. The number of infected neurons was on average 43% lower in microglia-containing triplecultures (mean count of infected cells = 12.50, 95% CI: 4.71 - 20.29, P = 0.16) when compared to infected co-cultures without microglia (mean = 21.80, 95% CI: 12.53 - 31.07) at 48 hpi. Tau phosphorylation at Ser202/Thr205 was increased 1.9- and 2.8- fold in infected triplecultures (Fig. 5E, F) at 48 hpi and 72 hpi, respectively, when compared to untreated controls. Taken together, the presence of microglia did not prevent neuronal infection by SARS-CoV-2 or infection-induced tau phosphorylation.

**Figure 5.**
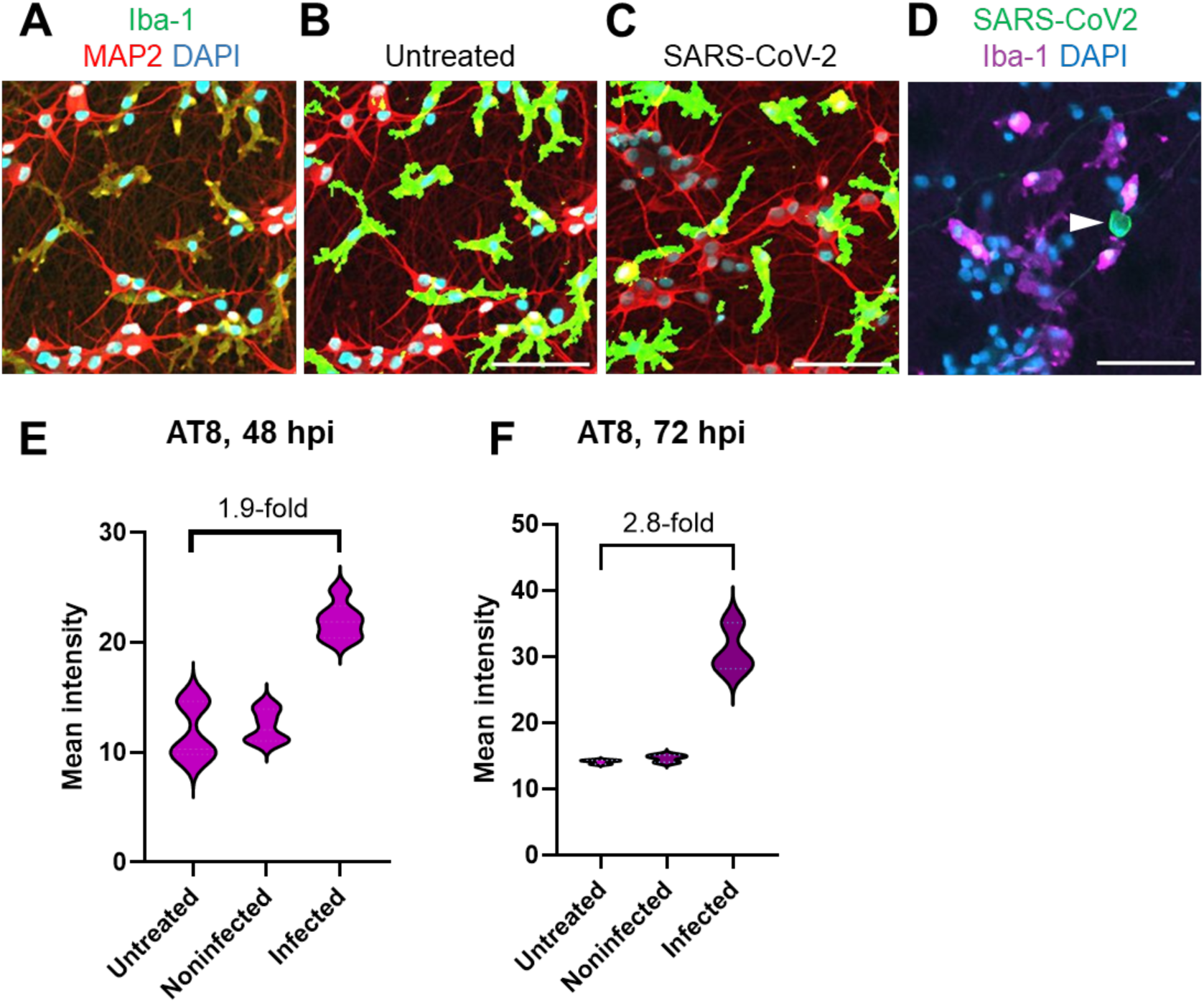
A. Representative image of anti-Iba-1 (microglia) and anti-MAP2 (neuron) staining of an untreated neuron-astrocyte-microglia co-culture. B. Same image but the Iba-1 staining has been thresholded with ImageJ into binary format to better display the microglial morphology. C. Similar thresholded image of SARS-CoV-2 infected culture. D Representative image of a SARS-CoV-2 infected neuron (stained with anti-N protein, arrowhead) among Iba-1 positive microglia in triplecultures. E. Mean intensity of AT8 fluorescent signal in untreated, treated but noninfected, and infected neurons at 48 hpi. N = 6 replicate wells. F. and at 72 hpi. N = 3 replicate wells. Scale bars 100 µm. All statistical significances were tested using Kruskal-Wallis test with Dunn’s correction for multiple testing. Only significant results are denoted. *, P ≤ 0.005.

## Discussion

### SARS-CoV-2 infection in iPSC-derived neurons is productive

Utilizing an ACE2-overexpression model, which resulted in a 2-10-fold increase in neuronal SARS-CoV-2 infection, we have demonstrated that endosomal cathepsins, particularly cathepsin-B, play a key role in SARS-CoV-2 entry in neurons. We also confirmed that infected neurons can produce and release infectious progeny virions. There has been a discrepancy in the literature as to whether neuronal infection leads to the production of new infectious virions (21,40,48–51), which could be due to differences in the experimental culture models and viral strains. In our model, viral RNA was released from neurons at least until 72h. However, the infectivity of the correspondent progeny virions peaked at 24 hpi and rapidly declined at later time points. These results could be explained by possible reduction in the stability of the released virions after 24 hpi, or by an antiviral response in the host cells which dampens the infectiousness of the virions. Further experiments will be required to test these possibilities.

### SARS-CoV-2 neuronal infection and the usefulness of cathepsin inhibitors

Based on available evidence, SARS-CoV-2 has multiple entry routes into the cell and the employed entry mechanism is dependent on the availability of entry factors and proteases that cleave the viral spike allowing fusion (37). In cells lacking suitable surface proteases, such as TMPRSS2, the virus must enter the cell through the endo-lysosomal compartment where the spike protein is cleaved by intralumenal proteases, such as cathepsins, to release the viral single-stranded RNA into the cytosol (12).

Previously, Zhao et. al. have highlighted the importance of cathepsin L in endo- lysosomal SARS-CoV-2 entry. They reported that SARS-CoV-2 infection increased cathepsin L expression in immortalized human cell lines *in vitro* further increasing the infection rate. *In vivo*, higher plasma cathepsin L levels correlated with more severe disease type while cathepsin B levels did not (26). In contrast, Hashimoto et. al. reported that the cathepsin B inhibitor CA-074-ME, and a *CTSB* knock-out.were effective in blocking SARS-CoV-2 infection in hiPSCs, particularly when used in combination with inhibitors of TMPRSS2 (52).

In our experiments, the small molecule inhibitor CA-074-ME, which has the strongest affinity for cathepsin B, efficiently blocked neuronal SARS-CoV-2 infection while the cathepsin L inhibitor SB412515 was less effective. In Vero E6 cells, where the infection of SARS-CoV-2 has been shown to depend mainly on cathepsin L, the small molecule SB412515 was more effective that the CA-074-ME. The stronger effect of CA-074-me compared to SB412515 at low concentrations suggests that, in this neuronal model, cathepsin B is more important for cleaving SARS-CoV-2 spike protein and initiating cell fusion than cathepsin L. Supporting this result, the neurons used in our co-culture model express no TMPRSS2, low levels of ACE2 (12), and twofold higher levels of cathepsin B compared to cathepsin L in mRNA level. In addition, CA-074-ME also showed a strong inhibitory effect in genetically non- modified cortical brain organoids, where the expression levels of neuronal cathepsin B are higher compared to cathepsin L (30). Furthermore, it has been reported that in vivo human excitatory neurons express higher levels of cathepsin B and L than ACE2 or TMPRSS2, with cathepsin B levels being, on average, four times higher than cathepsin L (53).

Cathepsin inhibitors are never fully specific, and they can inhibit other cathepsins when the concentration is high enough (54,55). While a relatively low specificity of the drugs would hamper mechanistic studies, it could be an advantage for possible use *in vivo*, where the residual pro-viral activity of other cathepsins would be inhibited by moderate doses of the same inhibitor. Pre-clinical studies in rodent models confirmed that CA-074-ME is well tolerated (41,56,57).Taken together, our results suggest that small molecule CA-074-ME is a promising drug candidate to prevent the potential SARS-CoV-2 spreading into the nervous system.

In our study, the cathepsin L inhibitor SB412515 was more effective against the Omicron XBB.1.5 variant compared to the parental Wuhan strain.

The utilization of different entry routes and different cathepsins by SARS-CoV-2 may change as the virus mutates. So far, it has been reported that the Omicron variants do not efficiently use TMPRSS2 but rather rely on other cell surface proteases or endosomal cathepsins (52,58,59). The identification of ‘pan-variant’ antivirals, such as the CA-074-ME, will aid in the development of therapeutic interventions that could limit the risk of neuronal invasion by the evolving SARS-COV-2.

### Cellular stress and signs of neurodegenerative diseases after SARS-CoV-2 infection

We observed a time dependent increase in the levels of HIF-1α in infected neurons. HIF-1ɑ is a constitutively expressed transcription factor that is rapidly degraded by the ubiquitin-proteasome pathway during physiological non-stressed conditions. During hypoxia, the protein’s stability is increased, leading to increase in its cytoplasmic concentration (60). HIF-1ɑ stabilization leads to changes in cellular metabolism, mainly induction of glycolysis, which mitigates the harmful effects of hypoxia (61). This protein has been previously associated with other viral infections such as influenza A (62), cytomegalovirus (63,64), and the human immunodeficiency virus (HIV) (65), and SARS-CoV-2 (66–70). Accumulation of HIF-1α has been previously shown in proximity with SARS-CoV-2 infected cells in a 3D organoid model (10). The translocation of HIF-1ɑ into the nucleus can trigger the antiviral interferon response and cytokine production (65,70,71). It may protect cells from infection (64,69,72) but also exacerbate tissue damage stemming from an over- activated immune system. On the other hand, HIF-1ɑ accumulation and associated metabolic changes have been reported to enhance viral replication and entry into cells (65,70,73,74). Taken together, it remains to be determined whether HIF-1ɑ accumulation is more beneficial to the host cell or the virus.

At 48 hpi, we also detected increased levels of MIF, MCP-1/CCL2, Serpin E1 and CXCL12 in neuron-astrocyte co-cultures following SARS-CoV-2 infection. These cytokines/chemokines promote the migration of immune cells (75–79), are increased in the plasma and/or serum of COVID-19 patients, particularly in severe cases (80–84) and are known to be regulated by HIF-1ɑ (85–88). There is some indication that endothelial plasminogen activator inhibitor (Serpin E1) can limit SARS-CoV-2 spreading (89,90). The high expression MIF allele has been associated with reduced susceptibility to symptomatic SARS-CoV-2 but increased disease severity in the symptomatic patients (91). Thus, MIF, MCP-1/CCL2, Serpin E1 and CXCL12 are likely to be a part of the anti-viral response mounted by neuron-astrocyte co-cultures and could be involved, at least in part, in the inhibition of virus infectivity observed in our model system after 24 hpi.

We observed a 43% reduction in the number of infected cells in microglia-containing triplecultures compared to cultures without microglia. An earlier report from a human ACE2 knock-in transgenic mouse model has shown that microglia promote the inflammatory response but do not restrict the infection (92). Interestingly, increased Serpin E1 levels have been reported to reduce microglial phagocytosis of zymosan-coated particles (75).

Reduction in the number of infected neurons over time suggests that the neurons do eventually succumb to the infection in this model system. However, we were unable to detect other signs of viral infection-induced cell death using multiple biochemical techniques. This is likely caused by the facts that 1) our co-cultures exhibit some cell death also under normal conditions; that 2) the level of viral infection was low, and that 3) it was difficult to catch the right moment to do co-staining for SARS-CoV-2 infection and cell death markers.

We also monitored the possible interactions between SARS-CoV-2 and AD-associated aggregating proteins. The SARS-CoV-2 spike protein has been reported to have amyloid-like properties through which it might be able to interact with other amyloid-like proteins (93). According to Eberle et. al. the SARS-CoV-2 3CL protease (3CLpro) induces tau aggregation *in vitro* (94). Consistently, tau accumulation has been reported in i*n vitro* models of neuronal SARS-CoV-2 infection, including hiPSC-derived brain organoids (40) and cortical-blood vessel assembloids (10), SH-SY5Y neuroblastoma cells (95), and in the brain of mice overexpressing human ACE2 (95). Here, we have detected tau phosphorylation at S202 / T205 (AT8) and mislocalization from the neurites to the soma of SARS-CoV-2 infected neurons in 2D neuron-astrocyte co-cultures and microglia-containing triplecultures.

According to Neddens et. al., increased tau phosphorylation at S202/T205 only appears at Braak stages V and VI (a semiquantitative measure of the level of tau pathology), which suggests that this change is associated with late-stage tau pathology (96). Consistent with our result, they also report that phosphorylation at S396 is generally weak with only minor progression (96). Our results, thus, indicate that SARS-CoV-2 induced neurodegeneration-like changes in infected neurons.

## Conclusion

This study demonstrates that SARS-CoV-2 can infect iPSC-derived neurons, and that endo-lysosomal cathepsins, particularly cathepsin B, play a crucial role in viral entry. Infected neurons can produce and release infectious virions, the infectivity of which peaks at 24 hours post-infection. The study also notes signs of cellular stress, the activation of antiviral response, and increased tau phosphorylation in the infected neurons suggesting potential links to neurodegenerative processes. The research highlights cathepsin inhibitors CA-074-ME and SB412515 as potential treatments for SARS-CoV-2 infections in the brain.

## List of abbreviations

ACE2: angiotensin-converting enzyme 2
CXCL12: CXC motif chemokine ligand 12
DAPI: 4’,6-diamidino-2-phenylindole (nuclear marker)
GFAP: glial fibrillary acidic protein
HIF-1α: hypoxia-inducible factor 1 α
hIPSC: human induced pluripotent stem cell
hpi: hours post-infection
Iba-1: ionized calcium-binding adapter molecule 1 (microglia marker)
IL-6: interleukin 6
IL-8: interleukin 8
IP-10: interferon gamma-induced protein 10
MAP2: microtubule-associated protein 2 (neuronal marker)
MCP-1: monocyte chemoattractant protein-1
MIF: macrophage migration inhibitory factor
MOI: multiplicity of infection
qRT-PCR: quantitative real-time polymerase chain reaction
SARS-CoV-2: severe acute respiratory syndrome coronavirus 2 Tau microtubule-associated protein tau
TMPRSS2: transmembrane serine protease 2

## Materials and Methods

### Immortalized cell line cultures

We cultured the African Green monkey kidney cell line Vero E6, and the wildtype and ACE2 and TMPRSS2-overexpressing forms of the human alveolar basal epithelial cell line A549 (A549 wt and A549-AT) in 75 cm2 filter screw cell culture flasks (Greiner, Kremsmünster, Austria) in 12 mL of Dulbecco’s Modified Eagle’s Medium with high glucose (Sigma Aldrich, MO, USA) supplemented with 10% Fetal Bovine Serum (ThermoFisher Scientific), 1% MEM NEAA 100X (ThermoFisher Scientific, Waltham, MA, USA), 1% L-glutamine (XX), and 1% penicillin G-streptomycin (XX). Prior to the experiments, we passaged the cultures a few times using 0.25% Trypsin- EDTA (ThermoFisher Scientific). A day before the experiments, we seeded the cells into 96-well plates (PerkinElmer, MA, USA) at density 15 000 cells/well in 100 µL of culturing medium and kept at 37°C and 5% CO2.

### Generation and culturing of human iPSCs (hiPSCs)

During this study, we used two hiPSC lines (cell line 1 / MAD8 clone 1; cell line 2 / MAD1 clone 7) that have been previously characterized elsewhere (97). Briefly, we collected skin fibroblasts from healthy Finnish males with punch skin biopsies and reprogrammed into iPSCs using a CytoTune iPS 2.0 Sendai reprogramming kit (ThermoFisher Scientific). We expanded the selected hiPSC colonies on Matrigel (growth factor reduced; Corning, Corning, NY, USA) in Essential 8 medium (ThermoFisher Scientific) at 37°C, 5% CO2. We passaged the cultures with 0.5 mM EDTA approximately twice a week. We confirmed the pluripotent nature and genomic normality of the colonies by qRT-PCR, immunocytochemistry and karyotyping.

### hiPSC-derived ACE2-overexpressing NGN2-neurons

We have described the production and characterization of human NGN2-cortical neurons in Kettunen et. (13). Briefly, we transduced hiPSCs with lentiviruses containing the Tet-O-Ngn2-Puro and FUdeltaGW-rtTA constructs (both plasmids from Addgene; lentivirus packing and concentration by Alstem, Richmond, CA, USA) and expanded them to obtain frozen stocks. Then, we transduced the lines again using a pLenti-6.3-CMV-Blast-hACE2 lentivirus construct (98) and selected the transduced colonies using 1.5 μl/ml blasticidin for 48 hours. Afterward, we cultured the ACE2- NGN2-iPSC under normal culture conditions: in Essential 8 medium (ThermoFisher Scientific) on 35mm Matrigel (growth factor reduced, Corning)-coated dishes at 37°C and 5% CO2. We passaged the cells with 0.5 mM EDTA every 2-5 days.

For each batch of experiments, we initiated neuronal differentiation by adding 2 μg/mL doxycycline to E8 medium on a 60 to 70% confluent NGN2-iPSC plate (day 0). For the next three days, we cultured the emerging neuronal precursor cells (NPCs) with doxycycline and dual SMAD inhibitors (LDN-193189 and SB-431542B, Sigma Aldrich) in N2 medium (DMEM/F12 without l-glutamine, 1% GlutaMAX, 1% N2 [all from ThermoFisher Scientific] and 0.3% glucose). On day 2, we selected the NPCs by adding 5 μg/mL puromycin (ThermoFisher Scientific). On day 4, we plated the emerging neurons with astrocytes as described below.

### hiPSC-derived astrocytes

We have described the production and characterization of human astrocytes in Kettunen et. al. (13). Briefly, we first produced NPCs from human iPSCs by dual SMAD inhibition (10 μM SB-431542B and 200 μM LDN-193189 for 10 to 12 days) and expanded them on laminin-coated plates with 20 ng/mL bFGF for 2 to 4 days. Then, we moved the cells to ultra-low attachment plates (Corning) in astrocyte differentiation medium (DMEM/F12 without l-glutamine, with 1% GlutaMAX, 50 μM nonessential amino acids, 1% N2, 50 U/mL penicillin, 50 μg/mL streptomycin, 0.5 IU/mL heparin) supplemented with 10 ng/mL bFGF and epidermal growth factor. We cultured the resulting astrospheres for 6 to 10 months and cut them manually when necessary. For experiments, we dissociated the astrospheres with StemPro Accutase (ThermoFisher Scientific) for 10 min, triturated into a single-cell suspension, and plated on culture dishes as described below. For astrocyte monocultures, we plated the astrocytes on growth factor-reduced Matrigel (Corning) at a density of 50,000 cells/cm2 and maturated them in astrocyte differentiation medium supplemented with 10 ng/mL BMP-4 and CNTF (ThermoFisher Scientific) for 1 week.

### hiPSC-derived microglia

For microglia differentiation, we detached iPSC colonies with ReLeSR reagent (STEMCELL Technologies, Vancouver, Canada) and plated them at density 3-6 colonies per cm^2^ on Matrigel-coated 6-well plates (growth factor reduced, 1:200, Corning, Corning) in Essential 8 medium supplemented with 5 µM ROCK inhibitor Y- 27632. We then differentiated the cells into hematopoietic progenitors using the commercial STEMdiff Hematopoietic kit (STEMCELL Technologies) for 11 - 13 days. Then, we collected floating hematopoietic progenitors and plated them at density 7000-8000 cells per cm2 on Matrigel-coated 6-well plates. We cultured the cells for 12 - 14 day in microglial differentiation medium which contains DMEM/F12, 2× insulin-transferrin-selenite, 2× B27, 0.5× N2, 1× glutamax, 1× non-essential amino acids (all from ThermoFisher Scientific), 400 μM monothioglycerol (Merck, Rahway, NJ, USA), 5 μg/mL human insulin (Merck), 100 ng/mL recombinant human IL-34 (Sino Biological, Beijing, China), 50 ng/mL human TGF-β1, and 25 ng/mL human M- CSF (all cytokines from Peprotech, Thermo Fisher Scientific).

### hiPSC-derived neuron-astrocyte co-cultures and microglia-containing triplecultures

On day 4 of neuronal differentiation, we dissociated the neuronal precursor cells with Accutase StemPro Accutase (ThermoFisher Scientific) for 5 min and plated them with astrocytes on plates coated with 9 to 16 μg/cm2 poly-d-lysine and ∼1.5 μg/cm2 laminin (from a mouse Engelbreth-Holm-Swarm sarcoma; Sigma Aldrich). The densities we used are listed below. On day 7, we inhibited the proliferation of residual dividing cells with an overnight treatment with 10 μM floxuridine (Bio-Techne, Minneapolis, MN, USA). Since microglia are dividing cells, we added them to the triple-cultures only after the floxuridine treatment (earliest on day 10).

For culturing, we used a base medium of Neurobasal (ThermoFisher Scientific) supplemented with 1% GlutaMAX (ThermoFisher Scientific), 2% B27 without vitamin A (ThermoFisher Scientific), 50 μM nonessential amino acids (ThermoFisher Scientific), 0.3% glucose and 10 ng/mL glial cell line-derived neurotrophic factor (GDNF), brain-derived neurotrophic factor (BDNF). For neuron-astrocyte co-cultures we added also 10 ng/mL ciliary neurotrophic factor (CNTF) (ThermoFisher Scientific). For microglia-containing triplecultures we added 100 ng/mL IL-34 (Sino Biological) to support microglial maturation.

We used glass-bottom 96-well plates (cat. no. 6055302, PerkinElmer) for imaging studies, 35mm culture dishes for qRT-PCR, and 4-well plates for media collection. For imaging and qRT-PCR studies, we used the density 50,000 cells/cm^2^ for neurons and astrocytes, and 25,000 cells/cm^2^ for microglia. For experiments that required media collection, we used thick cultures with density ∼100,000 cells/cm^2^ for neurons and astrocytes and 50,000 cells/cm^2^ for microglia.

### hiPSC-derived cortical organoids with choroid plexus

The protocol and characterization of hiPSC-derived cortical organoids with choroid plexus (ChPCOs) has been previously published in Shaker et. al. (31). Briefly, we used human WTC hiPSCs (a gift from Professor Bruce Conklin) cultured on feeder- free system on Matrigel in mTeSR medium (STEMCELL Technologies, Vancouver, Canada). We initiated the organoid development by plating hiPSCs at density 20- 30% in TeSR1 medium on a 6-well plate coated with human embryonic stem cell (hESC)-qualified basement membrane matrix (STEMCELL Technologies). The next day, we induced the development of neuroectoderm by switching the medium to N2 medium (DMEM/F12, 2% B27, 1% N2, 1% MEM non-essential amino acids, 1% penicillin/streptomycin and 0.1% β-mercaptoethanol, all from ThermoFisher Scientific) supplemented with dual SMAD inhibitors (10 µM SB-431542 and 100 nM LDN-193189). This treatment was continued for three days with daily medium changes. On the fourth day, we detached the neuroectoderm colonies with 2.4 unit/ml dispase to induce the formation of neural spheroids. We cultured the spheroids in N2 medium for four days with a daily addition of the following supplements: 40 ng/ml bFGF (Biotechne, Minnesota, USA), 2 µM CHIR99021 (Sigma Aldrich) and 2 ng/ml BMP-4 (ThermoFisher Scientific). Then, we embedded the patterned neural spheroids in Matrigel (STEMCELL Technologies) and switched the medium to Neurobasal medium supplemented with 1% GlutaMAX, 0.5% N2, 1% B27, 0.25 µl/ml insulin (Sigma), 1% MEM non-essential amino acids, 1% penicillin/streptomycin, 0.35 µl/ml β-mercaptoethanol, 3 µM CHIR99021 and 5 ng/ml BMP-4. We cultured the organoids for two months with medium changes three times a week prior. All experiments were carried out in accordance with the ethical guidelines of the University of Queensland and with the approval by the University of Queensland Human Research Ethics Committee (Approval number-2019000159).

### Virus infections

We performed all infection experiments in a biosafety level 3 (BSL-3) facility following required university regulations. We used two SARS-CoV-2 Wuhan strains: The main strain was produced and harvested SARS-CoV-2 for the 2D co-culture and tripleculture experiments in a susceptible A549-AT cell line and stored them in aliquots in - 80°C until the time of the experiments (12). The virus genome was sequenced by deep sequencing at full genome coverage as described in (98), and the presence of an intact “furin cleavage site” in the spike gene was confirmed (sequence accession number MW718190). Alongside, we used an EGFP-tagged SARS-CoV-2 virus, which was a kind gift of Dr. Andres Merits (University of Tartu, Estonia)(99).

We infected 2D co-cultures and triplecultures with SARS-CoV-2 MOI 0.5 (and MOI 2.5 for the experiment shown in Fig. 1D). The base medium was Neurobasal (ThermoFisher Scientific) supplemented with 1% GlutaMAX (ThermoFisher Scientific), 2% B27 without vitamin A (ThermoFisher Scientific), 50 μM nonessential amino acids (ThermoFisher Scientific), 0.3% glucose and 10 ng/mL glial cell line- derived neurotrophic factor (GDNF), brain-derived neurotrophic factor (BDNF) and ciliary neurotrophic factor (CNTF) on day 35. In neuron-astrocyte-microglia triplecultures we replaced CNTF with 100 ng/mL IL-34. After infection, we cultured the cells at 37°C, 5% CO2 for up to 120 hours.

For the control experiments, we infected immortalized cell lines A549-AT and Vero E6 with the SARS-CoV-2 Wuhan strain in 96-well plates at MOI 0.06 and 0.03 (respectively). We used normal culture medium containing Dulbecco’s Modified Eagle’s Medium with high glucose (Sigma Aldrich, MO, USA) supplemented with 10% Fetal Bovine Serum (ThermoFisher Scientific), 1% MEM NEAA 100X (ThermoFisher Scientific, Waltham, MA, USA), 1% L-glutamine (Sigma Aldrich), and 1% penicillin G-streptomycin (Sigma Aldrich) as a base medium. After infection, we cultured the cells at 37°C, 5% CO2 for up to 120 hours.

For 3D organoid experiments, we used SARS-CoV-2 isolate QLD1517/2021 (alpha variant, GISAID accession EPI_ISL_944644) passage 2 (P2) which was kindly provided by the Queensland Health Forensic and Scientific Services, Queensland Department of Health. The isolate was amplified on Vero E6-TMPRSS2 cells to generate virus stock (P3) and confirmed by sequencing (17). Virus titers were determined by immuno-fluorescent foci-forming assay on Vero E6 cells (100) We infected the organoids with 10^6^ foci-forming units (FFU) of SARS-CoV-2 for 6 h at 37 °C. After incubation, we washed the organoids three times and cultured them in fresh media for 72 hours before processing them for immunohistochemistry

At the end of the experiments, prior to removal from BSL-3 facilities, we neutralized the virus by fixation by 4% PFA in PBS for 20 min (immunohistochemistry), or by the addition of 0.1% Triton X-100 in the cell culture medium (cytokine secretion assays). UV light was used as an additional inactivation step where it did not interfere with the analysis.

### Vero E6 reinfection assay

To assess the infectiousness of the virions produced by infected hiPSC neurons, we treated Vero E6 cultures with conditioned medium collected at 24, 48, 72, 120 hpi from 2D neuron-astrocyte co-cultures infected with the SARS-CoV-2 Wuhan strain. The Vero E6 cells were pre-seeded 24 hours earlier to 96-well plates at density 15 000 cells per well (four replicate wells per treatment). We treated them with a 1:1 mixture of conditioned medium and normal culture medium (DMEM with high glucose [Sigma], 2% fetal bovine serum [ThermoFisher Scientific], 1% MEM NEAA [ThermoFisher Scientific], 1% L-glutamine [Sigma Aldrich] and 1% penicillin G-streptomycin [Sigma Aldrich]) in a total volume of 50 μl. After two hours, we removed the conditioned medium and cultured the samples for 48 hours prior to fixation for immunocytochemistry using an antibody against SARS-CoV-2 N protein and the nuclear marker Hoechst. Finally, we analysed the samples using the high-throughput method described below.

### Drug treatments

To inhibit endosomal maturation, we treated 2D neuron-astrocyte co-cultures, and 3D cortical organoids with 0.2 µM apilimod dimesylate (Bio-Techne) and immortalized cell lines A549-AT and Vero E6 with 0.5 µM apilimod dimesylate. To inhibit direct SARS-CoV-2 spike protein cleavage and membrane fusion at the neuronal surface, we used 25 µM nafamostat mesylate (Bio-Techne). To inhibit the enzymatic activity of endo-lysosomal cathepsins B and L, we used 3, 10 and 30 µM CA-074 methyl ester (CA-074-ME, MedChemExpress) and 3, 10 and 22.5-30 µM SB412515 (Cayman Chemical), respectively. All drugs were dissolved in dimethyl sulfoxide (DMSO, Sigma Aldrich). Consistently, we used the addition of 0.1% DMSO as a control in wells without drug treatments. We performed all the drug treatment experiments on 2D neuron-astrocyte co-cultures using three to eight technical replicates in five independent batches. The drugs were added to the cultures on 96- well plates within one hour prior to the addition of the virus.

We performed the pharmacological inhibition experiments for the Wuhan strain in 2D neuron-astrocyte co-cultures in five independent batches using brain cells derived from two independent hiPSC lines. We used 3 to 6 replicate wells for each treatment / plate / experiment. For each repetition, we used different aliquots of virus and drug stocks. The experiment using the Omicron XBB.1.5 strain was done using six replicate wells per treatment.

### Immunocytochemistry for drug experiments and reinfection assay in 2D cultures

We fixed the cell cultures 20 min with 4% PFA in phosphate-buffered saline (PBS) and washed them two times for 3 min with washing buffer (Dulbecco’s PBS with 0.2 % BSA, pH 7.2). We performed quenching for 20 min with 50 nM ammonium chloride (Merck) in washing buffer followed by two times 3 min washes with washing buffer.

Then, we permeabilized the cells with 0.1% Triton-X-100 in PBS for 10 minutes and washed again. We diluted the primary antibodies (Table 2) in washing buffer and incubated overnight in 4°C. The next day, we washed the samples two times 3 min and applied a mixture of diluted secondary antibodies and nuclei staining (1:1000 dilution of Hoechst 33342) in washing buffer. We incubated the samples for one hour in room temperature protected from light, washed two times 3 min with washing buffer and finally replaced the washing buffer with PBS for imaging.

**Table 2:** List of primary and secondary antibodies.

| Antibody | Manufacturer | Conc. | Reference or catalog no. |
| --- | --- | --- | --- |
| Cleaved caspase-3 rabbit | Cell Signaling | 1:1,000 | 9661S |
| Cleaved caspase-3 rabbit | Cell Signaling | 1:500 | 9664 |
| GFAP rabbit | Agilent | 1:500 | Z033429-2 |
| HIF-1α mouse | Abcam | 1:500 | ab1 |
| Iba1 rabbit | Wako, VA, USA | 1:500 | 019-19741 |
| MAP2 chicken | Abcam | 1:500 | ab92434 |
| MAP2 guinea pig | Synaptic Systems,<br>Göttingen, Germany | 1:1000 | 188004 |
| Nucleocapsid rabbit | Jussi Hepoioki, University<br>of Helsinki | 1:2,000 | (101) |
| Tau, phosphorylated (AT8)<br>mouse | ThermoFisher Scientific | 1:500 | MN1020 |
| Tau, phosphorylated (S396)<br>mouse | Cell Signaling Technology | 1:500 | 9632 |
| Tubulin-3 mouse | PerkinElmer | 1:2,000 | 801201 |
| TTR sheep | BioRad | 1:500 | AHP1837 |
| Goat anti-guinea pig Alexa<br>Fluor 546 | ThermoFisher Scientific | 1:1000 | A-11074 |
| Goat anti-mouse Alexa Fluor<br>488 | ThermoFisher Scientific | 1:400 | A11001 |
| Goat anti-mouse Alexa Fluor<br>633 | ThermoFisher Scientific | 1:500 | A21052 |
| Goat anti-rabbit Alexa Fluor<br>488 | ThermoFisher Scientific | 1:1,000 | A11008 |
| Goat anti-rabbit Alexa Fluor<br>647 | ThermoFisher Scientific | 1:1000 | A32733 |
| Goat anti-chicken Alexa Fluor<br>568 | ThermoFisher Scientific | 1:1,000 | AB_2534098, A11041 |
| Donkey anti-rabbit Alexa Fluor<br>647 | ThermoFisher Scientific | 1:1,000 | A31573 |
| Donkey anti-mouse Alexa Fluor 555 | ThermoFisher Scientific | 1:1,000 | A32773 |
| Donkey anti-sheep Alexa Fluor | ThermoFisher Scientific | 1:1000 | A-11015 |

### Immunocytochemistry for co-culture characterization and visual assessment of HIF-1α and phosphorylated tau levels in 2D cultures

We first fixed the cell cultures with 4% paraformaldehyde and washed twice with PBS containing magnesium chloride and calcium chloride (termed PBS below). Then, we permeabilized the cells for 20 min with 0.25% Triton X-100 in PBS and blocked unspecific binding with 5% normal goat serum in PBS for one hour. We diluted the primary antibodies (Table 2) in 5% normal goat serum in PBS and incubated overnight at 4°C. The next day, we washed the samples three times for 10 min with PBS. We diluted the secondary antibodies 1:1,000 in 5% normal goat serum in PBS and incubated for one hour at room temperature. Finally, we washed the samples three times for 10 min with PBS and stained with the nuclear marker 4′,6-diamidino-2-phenylindole (dapi). The samples were imaged in PBS.

### Antibodies

A list of primary and secondary antibodies used in the study is provided in Table 2.

### Imaging for 2D co-cultures and triplecultures

We carried out high-throughput imaging for the drug experiments and reinfection assay using an ImageXpress Nano microscope (Molecular Devices, CA, USA) at the Light Microscopy Unit of the University of Helsinki. We used two Nikon objectives: 10× 0.3-numerical aperture (NA) Plan Fluor, WD 16 mm (pixel size, 0.655 μm) and 20× 0.45-NA S Plan Fluor ELWD, WD 8.2 to 6.9 mm (pixel size, 0.328 μm). For experiments using cell lines A549-AT and Vero E6 cells, we took nine images per well. For experiments using hiPSC neuron-astrocyte and neuron-astrocyte-microglia co-cultures, we took 16 images per well.

For the assessment of HIF-1α levels, tau phosphorylation status, and the calculation of the number of infected neurons at different time points, we used ZEISS Axio Observer microscope with Fluar 10x/0.50 M27 objective and Axio Observer.Z1 / 7 camera (all ZEISS, Oberkochen, Germany). We imaged the whole well using image tiling and stitched it into one image using the Zen Blue microscopy software (ZEISS).

### Image analysis of high-throughput images (2D cultures)

We carried out high-throughput image analysis using Cell Profiler 4.2.5 (102) to detect infected neurons and astrocytes and to measure the intensity of the fluorescent signal from antibodies against HIF-1α and anti-phosphorylated tau (AT8 and anti-S396). We analysed approximately 5000-10 000 cells per sample in three replicate wells for each treatment. HIF-1α expression was assessed using brain cells derived from hiPSC cell line 2. Tau phosphorylation was validated in both cell lines. First, we identified the nuclei using the dapi or Hoechst 33342 channel by the typical diameter of the nuclei and the Otsu thresholding method. Then, we expanded the nuclei by a few pixels to obtain the expanded nucleus, which served as a proxy for the neuronal soma. Then, we measured the anti-MAP2 intensity of these objects and classified the objects with intensity above a threshold as neurons. Objects with anti- MAP2 intensity below the set threshold were classified as astrocytes or other cells.

Then, we measured the intensity of the staining against SARS-CoV-2 N protein (anti-N) in the expanded nucleus. We considered cells with intensity above a defined threshold as infected. Alternatively, we detected infected cells manually based on anti-N staining and excluded them from the whole neuronal population to obtain infected and noninfected neuronal populations, for which we measured the mean intensity of target staining (anti-HIF-1α, AT8, anti-S396) (data shown in Fig. 3A, B, C, D, H, J and Fig. 4 C, D). In the control experiments using immortalized cell lines cells, we used a similar pipeline, but no cell type-specific markers were needed, as the cell line was homogeneous. Finally, we exported the data for analysis in Microsoft Excel or GraphPad Prism 9.0.

### Immunohistochemistry, imaging and quantification for 3D cortical organoids

We fixed the organoids in 4% PFA for 60 min at room temperature, washed with PBS three times for 10 min, and immersed in 30 % sucrose in PBS at 4°C. After the organoids had sunk to the bottom of the sucrose solution, we embedded them in a mixture of Optimal cutting temperature compound (O.C.T) and 30% sucrose solution at 3:2 ratio and froze on dry ice. We cut the samples into 14 µm thick sections and collected them on Superfrost slides (ThermoFisher Scientific). At the beginning of the staining protocol, we washed the samples three times for 10 min with PBS at room temperature. Then, we blocked unspecific binding for 1 hour with blocking solution containing 3% bovine serum albumin and 0.1% Triton X-100 in PBS followed by washes three times 10 min with PBS. We incubated the primary antibodies (Table 2) overnight at 4°C. The next day, we washed three times for 10 min with PBS and incubated with Alexa fluor-conjugated secondary antibodies (Table 2) for one hour in room temperature. Finally, we stained the nuclei with Hoechst 33342 (ThermoScientific).

We captured tiled z-stacks of the slides using a Zeiss AxioScan Z1 microscope and Zen 3.6 software. We then analysed the images using ImageJ as follows. First, we detected the nuclei using the Hoechst 33342 channel. Then, we detected cells as ‘objects’ using maximum entropy calculated as *S = -(sum)p*log2(p)* where p is the probability of a pixel greyscale value in the image. We overlaid the objects on images captured from different channels to calculate the percentage of infected MAP2 positive cells (infected neurons), and cells positive for both cleaved caspase-3 and MAP2 (dying neurons) for each treatment. We analyzed the data from three organoids per group using GraphPad Prism 9.0.

### Cytotoxicity assay for the used drugs

For the cytotoxicity assay, we treated Vero E6 and A549-AT cells with 0.5 µM apilimod, 25 µM nafamostat, 3, 10 and 30 µM CA-074-ME, or 3, 10 and 22.5 µM SB412515 for 24 hours. Since the drugs were dissolved in DMSO, we used the 0.1% DMSO as a negative control. We used 9 µM UCN-01 as a positive control since it is known to induce cell damage. We measured the cytotoxicity of these compounds using a CellTiter-Glo 2.0 cell viability assay (cat.no. G9241, Promega, Madison, WI,

USA) according to manufacturer’s instructions (document no. TM403, revised 1/23). Briefly, we acclimated the cells at room temperature for 30 min. Then, we added 100 μL of prewarmed (22 °C) CellTiter-Glo reagent (Promega) to the 100 μL of cell medium at room temperature. We lysed the cells for 2 min on an orbital shaker and left them on the bench covered from light for 10 min to stabilize the luminescence signal. We measured luminescence using a GloMax® Navigator Microplate Luminometer (Promega) with a 0.3 s integration time.

### Assessment of cytokine secretion

First, we collected samples of cell culture medium from infected and control neuron- astrocyte co-cultures as follows: We collected the medium from 4-well plates containing thick neuron-astrocyte co-cultures 48 hpi and stored them in -80°C until analysis. This medium was centrifuged for 15 min at 14 000xG prior to any downstream analysis. We then lysed the cell culture on the plate using RIPA buffer (ThermoFisher Scientific) supplemented with protease inhibitors (ThermoFisher Scientific) for 15 min on ice. We centrifuged the cell solution for 15 min at 14 000xG, collected the supernatant and stored in -80°C until we performed a bicinchoninic acid assay (BCA, ThermoFisher Scientific) to normalize the cytokine production to the protein concentration.

We screened the cytokine secretion profile from medium pooled from two replicate neuron-astrocyte co-culture wells per treatment using the Proteome Profiler Human Cytokine Array kit (Bio-Techne). The kit screens for 36 different human cytokines chemokines, and acute phase proteins using antibodies dot blotted on membrane. The complete list of targets is included in Sup. 4C.

After initial screening using Proteome Profiler Human Cytokine Array kit, we measured the secretion of interesting hits using the more quantitative Cytokine Bead Array kit (BD, New Jersey, USA) and the flow cytometer. Cytokines MCP-1, IP-10, MCP-1, IL-6, and IL-8 were selected based on our results and the availability of different flex kits from BD. We analysed the samples (two replicates per treatment) using BD Accuri C6 Plus flow cytometer with BD CSampler Plus software (BD Biosciences) located at the Biomedicum Flow Cytometry core unit, University of Helsinki. We used mean PE-Height fluorescence intensity values to construct the standard curves, and the concentration values were derived from standard curves using linear regression.

### qRT-PCR to assess RNA expression of ACE2, Cathepsin B, and Cathepsin L

We assessed the mRNA levels of ACE2, cathepsin B, and cathepsin L in hIPSC- derived wildtype and ACE2-overexpressing neurons, astrocytes, microglia, neuron- astrocyte co-cultures, and cell lines A549 wt (negative control) and A549-AT (positive control) using qRT-PCR. The brain cells were derived from hiPSCs from two independent donors. First, we isolated mRNA using a RNeasy minikit (Qiagen, Hilden, Germany) following the manufacturer’s instructions. We measured RNA concentration using NanoDrop. For cDNA conversion, we first diluted 500 ng of RNA in water and mixed it with random hexamer primer (ThermoFisher Scientific). Then, we incubated the samples 5 min at 65°C in a C1000 thermal cycler (Bio-Rad, Hercules, CA, USA). We added a synthesis mixture of 10 mM deoxynucleoside triphosphates, RNase inhibitor, and Maxima reverse transcriptase in reaction buffer (ThermoFisher Scientific) to the samples and run cDNA synthesis for 30 min at 50°C. We ran qRT-PCR using a Maxima probe/ROX qPCR master mix and the following TaqMan primers: ACE2 (HS01085333_m1), Cathepsin B (Hs00947439_m1), Cathepsin L (Hs00964650_m1), and GAPDH (Hs99999905_m1) (ThermoFisher Scientific) on a Bio-Rad CFX96 real-time system. We ran the samples at 95°C for 10 min followed by 40 cycles of 95°C for 15 s, 60°C for 30 s, and 72°C for 30 s. We normalized the results to human GAPDH expression using the Q-gene program (103).

### qRT-PCR to assess viral release

We harvested SARS-CoV-2 RNA by collecting 25 μL of cell culture medium at various time points post infection and stored in viral lysis buffer with RNA supplements (AVL buffer supplemented with 1% carrier RNA; Qiagen). We extracted RNA using a QIAamp viral RNA mini kit (Qiagen). We measured RNA concentration and quality using a NanoDrop 2000 spectrophotometer (ThermoFisher Scientific).

We run qRT-PCR in triplicated using reverse transcriptase-negative and template- negative controls and TaqMan Fast Virus 1-step MasterMix (ThermoFisher Scientific) (5 μL of the sample in a 20-μL total volume). The primers and probe we used for the reaction were ordered from Metabion: RdRP-SARSr-F2, 5′-GTG ARA TGG TCA TGT GTG GCG G-3′ as the forward primer; RdRP-SARSr-R2, 5′-CAR ATG TTA AAS ACA CTA TTA GCA TA-3 as the reverse primer; RdRP-SARSr-P2, 5′–6- carboxyfluorescein-CAG GTG GAA CCT CAT CAG GAG ATG C–BHQ-1–3′ as the probe. We ran qRT-PCR was with AHDiagnostics Agilent Technologies Stratagene Mx3005P using the following steps: reverse transcription for 5 min at 50°C, initial denaturation for 20 s at 95°C, and two amplification steps at 95°C for 3 s and 60°C for 30 s (the amplifications steps were repeated for 40 cycles).

### Statistical analysis

As per scientific convention, we performed statistical testing on relevant experiments. However, with small sample sizes, one should be generally careful of drawing statistical inferences. Since normal distributions are difficult to confirm with small sample sizes, we used the non-parametric Mann-Whitney U test for comparing two groups. For the comparison of more than two groups, we used the non-parametric Kruskal-Wallis test. When the main effect was statistically significant, we used post hoc multiple-comparisons tests. A cut-off point of P < 0.05 was used as a threshold for statistical significance. Data in figures are presented as means ± standard errors of the means (SEM). We performed all statistical analyses in GraphPad Prism 9.

## Declarations

### Ethics approval and consent to participate

Punch skin biopsies for hiPSC generation were collected from Finnish healthy males after informed consent. The study has received acceptance from the Research Ethics Committee of the Northern Savo Hospital District (license no. 123.13.02.00/2016).

All organoid experiments were conducted under the approval by the University of Queensland Human Research Ethics Committee (Approval number-2019000159).

### Consent for publication

Not applicable

### Availability of data and materials

Further information on methodologies and data can be provided upon request.

### Competing interests

We declare no competing interests.

### Funding

The study was supported by Helsinki Institute of Life Science, University of Helsinki, Finland (P.K, T.R, J.K); Academy of Finland (P.K, T.R, J.K); Biocenter of Finland (P.K, T.R, J.K); the Doctoral Programme in Microbiology and Biotechnology, University of Helsinki, Finland (R.O); the Strategic Research Council of Finland grant 335527 and 362468 (G.B); the Sigrid Juselius Foundation, Finland (G.B); the Faculty of Medicine, University of Helsinki, Finland (G.B); the Helsinki Institute of Life Science, University of Helsinki, Finland (G.B); Australian National Health and Medical Research council grant 2010917, Australia (G.B); the European Union Horizon Europe Research and Innovation Program grant 101057553 (G.B); the Instrumentarium Science Foundation Grant 240024 (R.O); the University of Queensland Amplify Fellowship (M.J). The funders had no role in study design, data collection and analysis, decision to publish, or preparation of the manuscript.

### Authors’ contributions

PK: design, data collection and analysis (2D cultures), interpretation, drafting of the article; JR: design, data collection and analysis (2D cultures), drafting of the article; TQ: data collection and analysis (2D cultures); RO: data collection (2D cultures); SE: data collection (organoids); MS: data collection (organoids); SM; image analysis (organoids), EW: guided the work, provided infrastructures and provided funding (organoids); MJ: guided the work, provided infrastructures and provided funding (organoids); JK: design, interpretation, drafting and commenting of the article, guided the work, provided infrastructures and provided funding; TR: design, data collection and analysis (2D cultures), interpretation, drafting and commenting of the article, guided the work; GB: design, analysis, interpretation, drafting and commenting of the article, guided the work, provided infrastructures and provided funding.

## Acknowledgements

We thank the Flow cytometry unit, Light Microscopy Unit, and the Neural iPSC core facility funded by Helsinki Institute of Life Science (University of Helsinki) and Biocenter Finland. We also thank Sanna Mäki and Tomas Strandin for their assistance for the PCR analysis to determine viral release. We are also grateful to Julian Sng (The University of Queensland, Australia) for helping with organoid infections.

## Additional material

File name: Supplementary figure 1

File format: Microsoft Word document (.docx)

Title of data: SARS-CoV-2 infection in hiPSC astrocytes

Description of data: Quantification of the infectivity of wildtype and ACE2- overexpressing hiPSC astrocytes by SARS-CoV-2 Wuhan strain.

File name: Supplementary figure 2

File format: Microsoft Word document (.docx)

Title of data: Cell death markers in SARS-CoV-2 infected co-cultures

Description of data: Representative images of fluorescent stainings of cell death markers in SARS-CoV-2 infected co-cultures.

File name: Supplementary figure 3

File format: Microsoft Word document (.docx)

Title of data: SARS-CoV-2 infection in Vero E6 and A549-AT cells

Description of data: Representative images of infection by SARS-CoV-2 Wuhan strain in Vero E6 and A549-AT cells at 24 hpi with or without drug treatments.

File name: Supplementary figure 4

File format: Microsoft Word document (.docx)

Title of data: Cytokine profiling of infected co-cultures

Description of data: Additional information on the cytokines and chemokines assessed by membrane assay (A) and quantification of IP-10 in cell culture medium by cytokine bead array (B).

## Notes

### Competing Interest Statement

The authors have declared no competing interest.

